# *DNA primase large subunit* is an essential plant gene for geminiviruses, putatively priming viral ss-DNA replication

**DOI:** 10.1101/2022.12.23.521785

**Authors:** Lampros Siskos, Maria Antoniou, Jose Riado, Montserrat Enciso, Carlos Garcia, Daniele Liberti, Danny Esselink, Andrey G. Baranovskiy, Tahir H. Tahirov, Richard G.F. Visser, Richard Kormelink, Yuling Bai, Henk J. Schouten

## Abstract

The family of *Geminiviridae* consists of more than 500 circular single-stranded (ss) DNA viral species that can infect numerous dicot and monocot plants. Geminiviruses replicate their genome in the nucleus of a plant cell, taking advantage of the host’s DNA replication machinery. For converting their DNA into double-stranded DNA, and subsequent replication, these viruses rely on host DNA polymerases. However, the priming of the very first step of this process, i.e. the conversion of incoming circular ssDNA into a dsDNA molecule, has remained elusive for almost 30 years. In this study, sequencing of melon (*Cucumis melo*) accession K18 carrying the Tomato leaf curl New Delhi virus (ToLCNDV) recessive resistance quantitative trait locus (QTL) in chromosome 11, and analyses of DNA sequence data from 100 melon genomes, showed a conservation of a shared mutation in the *DNA Primase Large subunit* (*PRiL*) of all accessions that exhibited resistance upon a challenge with ToLCNDV. Silencing of (native) *Nicotiana benthamiana PriL* and subsequent challenging with three different geminiviruses showed a severe reduction in titers of all three viruses, altogether emphasizing an important role of *PRiL* in geminiviral replication. A model is presented explaining the role of *PriL* during initiation of geminiviral DNA replication, i.e. as a regulatory subunit of primase that generates an RNA primer at the onset of DNA replication in analogy to *DNA Primase*-mediated initiation of DNA replication in all living organisms.

## Introduction

The family of *Geminiviridae* consists of more than 500 viral species which infect numerous monocot and dicot plants. Geminiviruses are divided into 14 genera based on genome organization, transmission vectors, and host range. The *Begomovirus* genus is the most diverse one and contains more than 400 whitefly transmitted species (Fondong et al., 2013; Fiallo-Olive et al., 2021), many of them being plant pathogenic, with a major economic importance in the agriculture of tropical and subtropical regions (Fiallo-Olive et al., 2020).

The genome of geminiviruses varies in size from ∼2.5 to 5 kb, and is organized in one (monopartite) or two (bipartite) circular single stranded (ss)-DNA molecules of approximately 2.5 kb, that are encapsidated into twinned icosahedral protein particles (Krupovic et al., 2009). Only begomoviruses include species with two genomic components, denoted DNA-A and DNA-B, which contain six respectively two open reading frames (ORFs). Proteins encoded by DNA-A are involved in genome replication (REn, replication enhancer protein; Rep, replication-associated protein; TrAP, transcriptional activator protein), particle assembly and insect transmission (CP, coat protein), control of gene expression, and modulation of host defense responses (AC2, AC4), whereas proteins encoded by DNA-B are required for intra-intercellular movement in host plants (MP, movement protein; NSP, nuclear shuttle protein) (Briddon et al 2010). Except for a sequence of approximately 200 nucleotides, called the common region, A and B components hardly share sequence homology. The common region contains various regulatory elements (e.g. iterons, TATA box) involved in replication and transcription, and a stem-loop structure that is essential for replication. The stem-loop structure contains a highly conserved nonanucleotide sequence (5’-TAATATTCA-3’) at its top, presenting the origin of viral replication, and is found in all geminiviruses (Bhattacharjee et al., 2022). Upon feeding of viruliferous whiteflies onto healthy plants, virions introduced into the host’s cytoplasm uncoat and release their ss-DNA, which traffics to the nucleus for replication (Kumar et al., 2019).

Geminiviruses replicate their genomes initially with the conversion of ss-DNA into a double stranded (ds) replicative intermediate (RI) that is used as template for rolling circle replication (RCR) which generates progeny single stranded DNA molecules (Bernardi et al., 2008). Geminiviruses do not encode DNA polymerases for replication of their DNA genome but rely on DNA polymerases encoded by the host. DNA polymerase α is responsible for generation of dsRIs by creating the complementary strand of ss-DNA whereas polymerase δ promotes subsequent accumulation of new geminiviral ss-DNA from the dsRI template (Wu et al., 2021). RCR is initiated by Rep with the introduction of a nick within the nonanucleotide sequence of a dsRI molecule (Ruhel et al., 2019). Apart from the essential functional role of *Rep* in replication initiation, *Rep* is also responsible for recruiting (Kushwaha, 2017) host factors involved in ubiquitination to promote trimethylation of histones on viral replisomes, leading to enhanced viral genome transcription. Viral genes are clock- and anti-clock wise oriented on the DNA genome and become expressed via bidirectional transcription of dsDNA molecules (Fondong et al., 2013).

Resistance to the geminivirus ToLCNDV has been investigated in cucurbits since the virus is a major threat to these crops. In melon (*Cucumis melo*) ToLCNDV resistance is attributed to two minor QTLs in chromosomes 2 and 12 and one major recessive QTL in chromosome 11: 30,112,563-30,725,907 nt (melonomics v.4) (Saez et al., 2017) whereas in *Cucurbita moschata* the same type of resistance was mapped in a major recessive QTL on chromosome 8 (Saez et al., 2020). This region of *C. moschata* is highly syntenic to chromosome 11 of melon, again at the similar (overlapping) locus ch11:30,045,251-31,108,850 nt. Within this syntenic region, two *C. moschata* genes (CmoCh08G001790, CmoCh08G001780) showed high impact mutations in the resistant parents, presumably leading to a knock-out of the gene, while another two (CmoCh08G001760, CmoCh08G001720) exhibited moderate changes that could alter the biochemical properties of the protein. The homologues of these genes in melon are MELO3C022313 (putative transmembrane protein), MELO3C022314 (putative F17L21.8), MELO3C022315 (MADS-box transcription factor 8-like) and MELO3C022319 (DNA primase large subunit), respectively. Due to the synteny of the QTLs between *C. moschata* and melon any changes in the aforementioned melon homologues could also be associated with resistance to ToLCNDV.

In our study, we evaluated these four candidate genes. Our plant genomic and functional data analyses revealed that plant DNA primase is essential for geminivirus pathogenicity. This enzyme is part of the DNA Primase complex, that synthesizes RNA primers for initiation of a DNA strand that is complementary to a single-stranded DNA template. A model is presented to explain the role of melon DNA primase in geminiviral ss-DNA replication.

## Materials and Methods

### Whole genome sequencing of K18 resistant melon line and SNPs detection

Leaf material from the ToLCNDV resistant line K18 carrying the recessive QTL in chromosome 11 and the susceptible line of similar genetic background K15 but lacking the ch11 QTL, was provided by Nunhems B.V in order to extract genomic DNA (gDNA) for whole genome sequencing (WGS). 200 mg of tissue per genotype were ground to fine powder using liquid nitrogen. The protocol of Healey et al. (2014) was followed for isolation of highly concentrated, high quality gDNA. 1.2 ug of non-contaminated, non-degraded gDNA, was sent to Novogene Cambridge Genomics Center, United Kingdom, for library preparation and WGS (PE150, Q30≥80%), using the Illumina Novaseq 6000 platform. Raw reads were aligned using Bwa mem with less stringent settings (-A1 -B1 -E1 -O1 -M) to reference genome Melon_v4.0, which includes the chloroplast and mitochondrial genome sequences. Duplicate reads were tagged using Mark Duplicates from Picardtools. Samtools was used for post-processing of the alignment files and Alfred for computing quality statistics of the alignment files (Danecek et al., 2021). Variants of chr 11 of both samples were joined called using FreeBayes (min-mapping-quality 2) and filtered using vcflib (vcffilter -f “QUAL > 1 & QUAL / AO > 10 & SAF > 0 & SAR > 0 & RPR > 1 & RPL >1”). SNPeff was used to predict the effect of protein-coding SNVs on the structural phenotype of proteins (De Baets et al., 2012).

### Virus induced gene silencing in *Nicotiana benthamiana* and viral inoculations

#### Plant material

115 *N. benthamiana* plants were grown at Unifarm, Wageningen University & Research. The greenhouse growth conditions were 24°C - 18°C day/night temperature with a photoperiod of 16 hours and 60% relative humidity (RH) for optimal germination and growth.

#### VIGS Constructs

A 321bp region of the *N. benthamiana DNA primase large subunit* (*NbPriL*) coding sequence targeting both gene transcript variants (*Niben101Scf00366g01013*.*1, Niben101Scf01950g03015*.*1*, Sol Genomics Network) was amplified through PCR with primers (ForwardNbPriL:5’-AGGATGCAACTTGGTCTATTTCTC-3’, ReverseNbPriL:5^’^-TTGTCTAACACATCCTCAACTGCT-3’). The region was chosen based on the best scoring of Sol Genomics Network VIGS tool for optimal virus induced gene silencing. The PCR product was purified and cloned into a pENTR™/D-TOPO™ vector with the cloning pENTR Directional TOPO® cloning kit. Using the Gateway™ LR Clonase™ Enzyme Mix the cloned insert was incorporated into the pTRV2 vector. *Agrobacterium tumefaciens* strain C58C1 was transformed using heat shock method with the pTRV2:NbPriL vector for downstream VIGS assays. In the same way the TRV:*GUS* and TRV:*PDS* constructs were previously generated.

#### Agroinfiltration with the VIGS construct

*Agrobacterium tumefaciens* cultures transformed with the various TRV2 constructs (TRV:Nb*PriL*, TRV:Nb*GUS* and TRV:Nb*PDS*) were mixed in 1:1 ratio with TRV1 cultures until a final OD_600_ of 2 was reached, and were eluted into MMA buffer (10 mM MES pH 5.6, 10 mM MgCl2, 200 Μm acetosyringone) (Norkunas et al., 2018). *GUS* encodes the beta-glucuronidase enzyme and is a known reporter gene that is not found in *N. benthamiana* genome (Hull et al., 1995). Two-week-old *N. benthamiana* cotyledons were syringe infiltrated with these mixtures. In plants with the TRV:*GUS* construct, the plant silencing machinery was turned on without any of the native plant genes being silenced. PriL silenced plants were compared to GUS silenced plants. Silencing of the *N. benthamiana PDS* gene was used as a positive control of silencing as successful silencing of this gene leads to a characteristic clearly visible developmental phenotype (photo bleaching). 40 plants were infiltrated with TRV:Nb*PriL*, 40 plants with TRV:Nb*GUS* and 5 with TRV:*NbPDSA. tumefaciens* cultures whereas the remaining 30 plants were kept wild type (WT).

#### Viral Inoculations

As soon as the *PDS* silenced plants exhibited the photo-bleaching phenotype (2 weeks post infiltration) showing successful silencing, agro-inoculation (Peyret et al., 2015) with various geminiviruses took place. Agrobacteria cultures (OD_600_=2) transformed with viral vectors of BCTV (Stanley et al., 1986), TYLCV (Morilla et al., 2005) or ToLCNDV (Lee et al., 2020) were eluted in MMA buffer and syringe infiltrated into 1-2 fully grown leaves of *N. benthamiana*. Eight plants were used per virus per VIGS construct, and six plants per virus for the WT controls. Eight TRV:*GUS*, eight TRV:*PriL* and 6 WT plants remained non-inoculated as controls. Viruses were maintained in different experimental blocks to prevent cross contamination, and the plants within each block were completely randomized.

### DNA and RNA extraction and quantification of viral titers and *PriL* transcripts

Young leaf tissue (non-infiltrated, non-inoculated) from all plants was harvested after the appearance of the first systemic viral infection symptoms. TYLCV inoculated plants were harvested 7 dpi as they started to exhibit symptoms earlier, whereas ToLCNDV and BCTV plants were harvested 10 dpi. The tissue was ground into fine powder using metal beads and Qiagen Tissue Lyser II Cat. No. / ID: 85300.

#### DNA extraction

Total DNA was extracted from the collected samples in order to determine viral DNA titers. For DNA extraction 200 mg of ground tissue per sample was mixed with 400 μl working buffer solution. The working buffer solution was prepared for 40 samples and contained: a) 12.5 ml extraction buffer (0.35 M Sorbitol, 0.1M Tris-HCl pH 8.0, 5 mM EDTA pH 8.0), b) 12.5 ml lysis buffer (0.2 M Tris-HCl pH 8.0, 0.5M EDTA pH 8, 2 M NaCl, 2% CTAB), c) 5ml sarcosyl 5% (w/v), d) 0.15g NaHSO_3_, e) 0.6g PVP-40. RNase A 20μg/ml was also added into the buffer mix working solution, and mixed. Samples were incubated at 65 °C for 1h. After incubation, two equal volumes of chloroform: isoamylalcohol (24:1) were added for extraction. After vortexing for 30s and subsequent centrifugation for 5 min at 13.000 rpm, the aqueous phase was collected and transferred to a clean microcentrifuge tube. An equal volume of cold isopropanol was added, and the tube inverted several times for DNA precipitation. DNA was pelleted by centrifugation for 5 min at 13.000 rpm. After isopropanol was discarded, DNA pellets were washed with 70% ethanol, and subsequently dried at room temperature for 15 min. DNA was resuspended in sterilized water and stored at -20C for further analysis.

#### RNA extraction

RNA was extracted from plants in order to determine transcription levels of Nb*PriL* after different treatments. RNA extraction was carried out using TRIzol according to the manufacturers’ protocol (Invitrogen). In brief, 1 ml of Trizol was added to each ground sample and the samples were vortexed for 30s. Next, 200 μL of chloroform was added, and samples again vortexed for 15s, followed by a centrifugation at 13.000 rpm for 30 min at 4 °C. From the aqueous phase 400 μL was transferred to a clean microcentrifuge tube and 400 μL of isopropanol was added and mixed. After incubation for 10 min at room temperature, RNA was pelleted during centrifugation at 13000 rpm for 20 min at 4 °C, and the pellet washed with 75% ethanol, air dried and subsequently resuspended in 50-70 μL of sterilized water. Thermofisher DNase I was used to remove (residual) DNA from the purified RNA samples. First strand cDNA synthesis was performed using BioRad iScript cDNA synthesis kit according to the manufacturers’ protocol.

#### Quantification of gene transcripts and viral titers

Quantitative real time PCR (qRT-PCR) was performed to determine gene expression levels and viral titers. Quantitative RT-PCRs were performed in technical triplicates, using a Bio-Rad CFX96 C1000 Thermal cycler with BioRad iQ SYBR Green supermix. Two sets of primers for the Nb*PriL* gene were used to quantify the total expression of the gene due to the two transcript variants *Niben101Scf00366g01013*.*1* and *Niben101Scf01950g03015*.*1*. Primers for viral titer quantification were designed based on the genomic sequence of each virus (Table 3) using total DNA extracted from the plants as input. Nb*PriL* total expression was calculated taking into account the CQ values of both transcript variants. *NbPriL* expression levels as well as viral titers were normalized to the housekeeping gene Nb*APR* (Solyc02g093150.2) (Liu et al., 2012). The 2-ΔΔCt method (Livak et al., 2001) was applied to assess relative transcript levels and viral titers. 2-ΔΔCt was calculated in Microsoft excel and t-test was used to discern the significant differences between control and test samples CQ values.

### Disease assays on melon accessions

Per CGN accession 12 plants were grown in a multispan greenhouse with passive ventilation through zenithal windows and cooling system for temperature control at SAKATA, Almeria, Spain. The same number of susceptible control “Coliseo” melon plants as well as the resistant control “GRAND RIADO” melon plants were added. Temperature range during the assay was 20-28 °C and relative humidity 60-90%. Each plant genotype was maintained in a different experimental block. Whiteflies collected from zucchini plants infected with ToLCNDV were released onto 1 week old plants (5-10 adults/plant). 20 days post inoculation access of viruliferous whiteflies, plants were scored on disease severity. Taking into account the systemic symptoms of ToLCNDV in melon as well as the overall fitness of the plant, a scale of 1-10 of disease score was used (1: fully asymptomatic – 10: fully symptomatic).

### 3-D modelling of PriL-PriS protein models

The coordinates of the melon PriS and PriL three-dimensional models were obtained from the AlphaFold database (accession codes A0A5A7T8Y3 and A0A1S3CBL6, respectively) (Jumper et al., 2021). Then melon PriS, PriLNTD and PriLCTD, domains were individually fitted into the model of human primase initiation complex (Baranovskiy et al., 2016) using the “align to molecule” function in PyMOL (The PyMOL Molecular Graphics System, Version 2.0 Schrödinger, LLC.). The linker between PriLNTD and PriLCTD was rebuild and refined using Coot (Coot3Emsley et al., 2010). The sites for the binding of metal ions and substrates are well conserved between the melon and human primases, providing their straightforward transfer from the human to the melon model.

## Results

### Sequencing of a ToLCNDV resistant melon source shows four changes in *DNA primase large* subunit coding sequence

Plant material of melon accession K18 carrying the ToLCNDV recessive resistance QTL was sequenced in order to examine potential changes in the four top candidate gene coding sequences compared to the susceptible reference. No high impact changes that could result into early stop codons and gene knock outs were found in any of the four candidate genes examined in K18. A more detailed look revealed no changes whatsoever in the coding sequence of the first candidate i.e. MELO3C022313 compared to the susceptible reference genome. The coding sequence of the second candidate, MELO3C022314, showed two SNPs, one synonymous *T-C* 30,276,085 nt and one moderate impact SNP *G-C* 30,275,808 resulting into a conserved *L-V* aa change in position 100 of the protein. A high impact, frameshift mutation (TA insertion, ch11: 32734434, v.3.6) that could result to a knock-out protein was previously found in MELO3C022314 of CGN24928 accession. However, subsequent disease assays showed that plants of this accession carrying the frameshift mutation in MELO3C022314 were susceptible to ToLCNDV, indicating that this gene is unlikely the susceptibility gene. As a potentially loss of function MELO3C022314 protein could not confer resistance to the virus, this gene was excluded as candidate susceptibility gene. The coding sequence of the third candidate *MELO3C022315* revealed only one synonymous nucleotide substitution (*A-G*, ch11: 30,297,365 nt) which does not impact the encoded protein sequence. However, and interestingly, five changes were found in the coding sequence of the last candidate gene, *MELO3C022319*. One of the nucleotide substitutions was synonymous, having no effect on the produced protein, whereas the other four resulted into amino acid changes in the *DNA primase large subunit* (PriL) protein (Table 1). Taken altogether, changes in the *DNA primase large subunit* most likely caused for the resistance mapped in chromosome 11.

**Table 1.** Overview on the four types of changes observed in the amino acid sequence of DNA Primase Large subunit (PriL) of melon accession K18, carrying the ToLCNDV recessive resistance QTL on chromosome 11. We used melonomics genome version 4.0.

| Change | SNP / genomic position | Amino acid change / position in protein | Polarity change |
| --- | --- | --- | --- |
| A | *T-C* / 30,355,908 | *Y-H* / 4 | Polar to basic polar |
| B | *C-T* / 30,354,702 | *P-L* / 125 | Polar to non-polar |
| C | *T-G* / 30,354,543 | *S-R* / 143 | Polar to basic polar |
| D | *C-A* / 30,354,303 | *A-E* / 165 | Non-polar to acidic polar |

### Silencing *PriL* resulted in reduced viral titers of ToLCNDV and two other geminiviruses in *Nicotiana benthamiana*

We knocked down the *PriL* homologue (Nb*PriL*) by virus induced gene silencing vector in *Nicotiana benthamiana*, and subsequently we challenged the plants with ToLCNDV. In order to investigate the involvement of PriL in the susceptibility to geminiviruses at family and genus level two other distinct geminiviruses were included: *Tomato yellow leaf curl virus* (TYLCV) (*Geminiviridae*: begomovirus) and *Beet curly top virus* (BCTV) (*Geminiviridae:* curtovirus). *N. benthamiana* plants verified for a reduction in the level of Nb*PriL* transcripts exhibited a characteristic phenotype that appeared 8-9 days post infiltration with the VIGS construct. This phenotype was consistently observed in all replicates and included shorter internodes and stunting that persisted throughout the experiment. Despite the stunted phenotype of plants infiltrated with TRV:Nb*PriL*, when compared with plants infiltrated with a TRV:Nb*GUS* negative control construct, the former plants were able to survive without any further defects (Figure1a).

**Figure 1.**
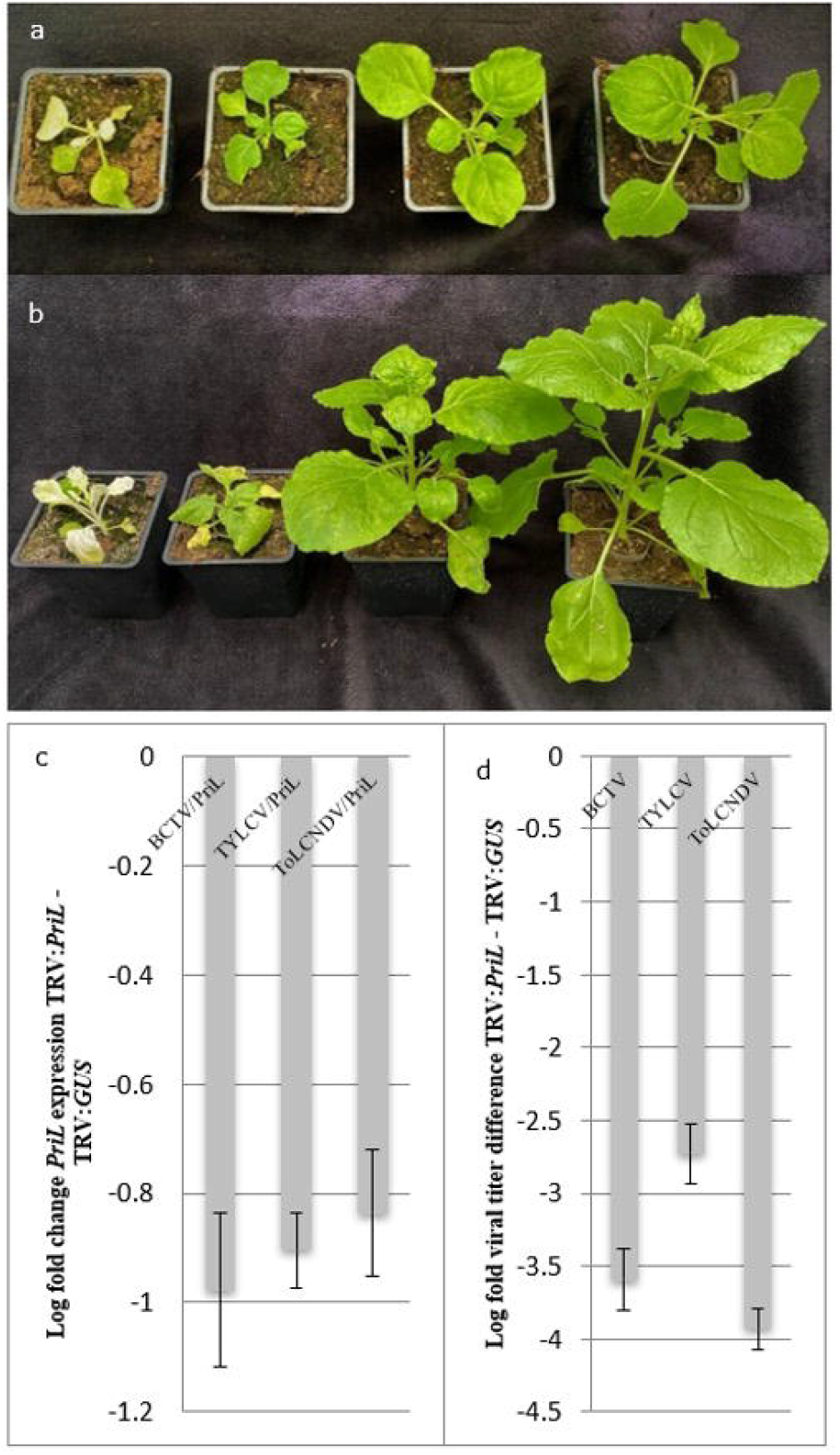
**a**. *Nicotiana benthamiana* plant phenotypes 14 days post infiltration with VIGS constructs. From left to right: TRV: Nb*PDS*, TRV: Nb*PriL*, TRV: *GUS*, Wild Type (WT); **b**. ToLCNDV inoculated *N. benthamiana* plant phenotypes 10 days post inoculation. From left to right: TRV: Nb*PDS*, TRV: Nb*PriL*, TRV: *GUS*, Wild Type (WT). Inoculation with the geminiviruses TYLCV and BCTV led to similar phenotypes compared to ToLCNDV. **c**. Average log fold *PriL* expression change of TRV:*PriL* plants, inoculated with the different viruses, compared to TRV: *GUS* plants. **d**. Average log fold difference in viral titers between TRV: *PriL* and TRV: *GUS* plants. Significant difference between TRV:*PriL* and TRV:*GUS* (p < 0.05).

Two weeks post infiltration with the VIGS constructs, *N. benthamiana* plants where agro-inoculated with infectious clones of ToLCNDV, TYLCV and BCTV. In plants that had received the silencing construct *TRV-NbGUS* disease symptoms started to show up one week post inoculation with TYLCV, consisting of leaf curling and light yellowing of the upper leaves. Symptoms were more prominent in TRV:Nb*GUS* infiltrated plants than in non-silenced wild type *N. benthamiana*. Three days following the symptoms in TYLCV challenged plants, first symptoms were observed in ToLCNDV (Figure1b) and BCTV challenged plants, and similarly revealed mild leaf curling and yellowing of the upper leaves, especially in the TRV:*GUS* plants. However, and in contrast, during this entire period of monitoring no symptoms appeared upon a challenge with any of the geminiviruses in plants silenced on *PriL* by the TRV-Nb*PriL* construct.

Upon the onset of the first viral symptoms in TRV:*GUS* plants (i.e. 7 dpi for TYLCV and 10 dpi in ToLCNDV and BCTV), expression levels of Nb*PriL* and viral titers were determined. As expected, *PriL* expression levels in plants that had received TRV:*PriL* were significantly reduced compared to the control plants containing TRV:*GUS* (Figure 1c) and WT plants (p <0.05). When viral titers were determined, significantly lower titers were observed for all viruses in TRV:*PriL* treated plants compared to the negative control, TRV:*GUS* treated plants (p<0.05). A further look showed that in *PriL*-silenced plants TYLCV viral titers were higher than the other two viruses (p<0.05). Altogether, silencing of *PriL* led to significant reduction in viral titers of all three geminiviruses, strengthening its role as a susceptibility factor for geminiviruses.

### DNA sequence data of 100 melon genomes show a conservation of *PriL* alleles with a shared mutation among accessions with ToLCNDV resistance

Having identified four mutations in *PriL*, leading to an amino acid substitution in the resistant melon parent K18, their occurrence was analyzed in the *PRiL* gene of 100 melon accessions, as sequenced by Demirci et al. (2021). In addition, ToLCNDV disease assays were performed on those accessions to investigate whether, and if so which mutations conferred ToLCNDV resistance, similarly to the K18 accession. ToLCNDV disease assay data were collected from greenhouse screenings in Spain using whitefly inoculations. In 18 out of the 100 accessions one or more mutations were present in *PriL* that were also observed in K18, and leading to an amino acid substitution. Those alleles contained all the mutations as present in K18 (ABCD) or only some of them (ACD, AB, C, D, B, BD, BCD) (Table 1, Supplementary table 2).

From all four mentioned amino acids changes only mutation A consistently associated with ToLCNDV highly resistant phenotypes (symptomless). All evaluated accessions that harboured the Y to H substitution on position 4 (A) of *PriL* appeared to exhibit resistant phenotypes to ToLCNDV comparable to the resistant control genotype. Two accessions were carrying the same aa change according to sequencing data, however on a heterozygous state, and appeared to be susceptible to ToLCNDV. One accession carried mutation A in a homozygous state but only about half of the plants that were phenotyped exhibited highly resistant phenotypes to the virus. The rest of the CGN140853 plants were exhibiting fully or intermediately resistant phenotype scores (D.I. with 10 = fully symptomatic and 1 = fully symptomatic: 7.0 in 3/18 plants, 5.0 in 3/18 plants and 3.0 in 2/18 plants, 1.0 which means fully asymptomatic in 10/18 plants). Mutation B exhibited a correlation with phenotypes of intermediate resistance, independently of the presence of mutation A (Supplementary Table 1).

From these observations we conclude that the amino acid substitutions from tyrosine Y to histidine (Y to H) at position 4, and from proline to leucine (P to L) at position 125 are the most likely changes in PriL in the resistant genotype K18, leading to loss of its function for the virus, but maintenance of the protein’s function for the host plant.

## Discussion

The investigation into the genetic cause of recessive resistance to ToLCNDV on chromosome 11 of melon identified the *DNA primase large subunit* (*PRiL*) gene as the most likely candidate. We found non-synonymous mutations in the *PRiL* gene in the resistant K18 melon line, which were also present in multiple other resistant melon accessions with diverse genotypes. The resistance of these accessions was determined through disease index scoring. In future studies, the quantification of viral titers could be included to confirm that the observed resistance is reflected in reduced viral titers.

To investigate the functional role of *PRIL* in ToLCNDV susceptibility a VIGS approach was implemented in *N. benthamiana* (Kumar et al., 2014) due to the inefficiency of the method in melon. Efficient silencing of the native *PriL* homologue in *N. benthamianana* resulted into characteristic developmental phenotypes but also into significantly reduced viral titers for all evaluated geminiviruses ToLCNDV, TYLCV and BCTV. In future experiments the effect of *PriL*-silencing will be extended to other viruses (e.g RNA viruses) that do not rely on PriL as a susceptibility factor for their multiplication, to further substantiate the findings from this study. Our findings indicated a high correlation of melon’s PriL A-type mutation (involving an Y-H substitution, Table 1) with ToLCNDV resistant phenotypes. There was one exception to this correlation, which was observed with accession CGN140853. Although the plants in this accession were homozygous for the A-type mutation, they were still somewhat susceptible to ToLCNDV, although their susceptibility was reduced compared to the susceptible control. Since resistance against ToLCNDV in melon is known to be controlled by more than one QTL (Saez et al., 2017), it could be that the minor QTLs in chromosome 2 and 12, with an additive effect on resistance, are absent in this particular accession resulting into a more susceptible phenotype. Moreover, the resistance in chromosome 11 could be the result of more than the A-type mutation in PriL. Type B mutation (Table 1) could be another factor of resistant phenotypes. The B mutation gave indications of conferring intermediate resistance (independently of the A presence) in CGN140808 and CGN140870, but the correlation of the mutation with the effect was weaker. Altogether, the results indicate that *PriL* plays a major role in the pathogenicity of ToLCNDV and other geminiviruses in melon. Eventually, at least two aminoacid changes can alter *PriL* in a way that still supports the protein’s role for the plant but hamper viral replication.

The plant replication machinery includes several essential genes that are required for DNA replication of the plant. Apart from DNA polymerases, DNA primases are also essential components for DNA replication of the plant. Their role is to synthesize RNA primers that polymerases subsequently elongate on both DNA template strands of the replication fork for the generation of new dsDNA, since polymerases alone are unable for *de novo* dsDNA synthesis (Frick et al., 2001, Kuchta et al., 2010). Eukaryotic primases are heterodimers consisting of two distinct subunits: a small one, DNA Primase Small subunit (PriS), which carries the active site for RNA primer (∼9 nt) synthesis, and a large one, PriL, which determines the activity of the primase and the transfer of the synthesized primer to Polymerase α (Polα) (Baranovskiy et al., 2016). In organisms with dsDNA, primer synthesis happens only once in the continuous leading strand replication, which becomes extended at its 3’ terminus by Polα and Polε for the formation of the complementary strand. In the lagging strand the orientation is opposite. Consequently, RNA primers need to be newly generated constantly, with following extension by Polα and Polδ leading to Okazaki fragments. With this discontinuous strand replication, sequential Okazaki fragments altogether generate the complementary strand of the lagging strand (Zhou et al., 2019).

Considering the fundamental role of *PriL* in cell replication, the developmental phenotype that was observed after silencing of Nb*PriL* was not unexpected since plant DNA replication was likely inhibited to some extent. Although the development of a normal phenotype was compromised after silencing of Nb*PriL*, the residual expression levels (∼50% transcript availability) of Nb*PriL* were still enough to maintain viability and functionality of the plants throughout the experiment. Another indication of the significance of *PriL* in plant viability was the absence of high impact mutations (e.g coding sequence deletions, insertions) that could lead to a knock-out of the protein in any of the 100 melon genotypes examined. Normally, plants have only one gene copy of *PRiL*. Loss of function of that gene would be lethal, explaining the absence of high impact mutations in *PriL* in the 100 melon genomes. This implies that only subtle mutations in *PriL* would be allowed, such as amino acid substitutions. This is in agreement with our findings.

Remarkably, despite the thorough studies (Wu et al., 2021) on the role of host polymerases in geminiviral replication there were still no suggestions, comments or experimental data that refer to a possible involvement of host encoded DNA primases in replication of viral DNA. The only study that gets close to this point goes back 30 years ago: Saunders et al. (1992) observed that the conversion of ss-into dsDNA of the geminivirus African cassava mosaic virus (ACMV) is initiates in a region very close to the hairpin loop with the conserved nonanucleotide sequence. There, a small RNA nucleotide sequence is preceding the complementary DNA strand, at the start of this complementary strand. This observation pointed to the possibility of priming with a RNA oligonucleotide, of which the origin and generation have remained elusive so far.

Based on findings from this study it is now tempting to hypothesize that *PriL* together with *PriS* from the host generates an RNA oligonucleotide as the very first step to prime the complementary strand on the single stranded DNA of the virus, thus enabling the conversion of incoming ssDNA into dsDNA (Figure 2). Whereas the ‘natural’ location of the DNA primase in living organisms is close to the replication fork, the single-stranded DNA geminiviruses do not have such replication fork. However, all geminivirus DNA components contain a hairpin loop in the common region with the conserved nonanucleotide sequence “TAATATTAC” at its top (Figure 2), which resembles a replication fork of DNA replication complexes (Heyraud et al., 1993). The base of this hairpin loop could present the initiation site for DNA Primase to start RNA oligomerization. Docking of the host DNA primase just upstream of the hairpin loop would be in agreement with the findings from Saunders et al. (1992), who mapped an RNA primer at this site. Since all Geminiviruses need the conversion from incoming single-stranded to double-stranded DNA, and all geminiviruses contain the structurally highly conserved region with a hairpin loop, it is likely that priming by DNA primase at the hairpin structure is generic to all geminiviruses, irrespective of the host. This also turns PriL into a susceptibility factor for geminiviruses that do not limit to melon, and opens the possibility for its exploitation in other important crops as well.

**Figure 2.**
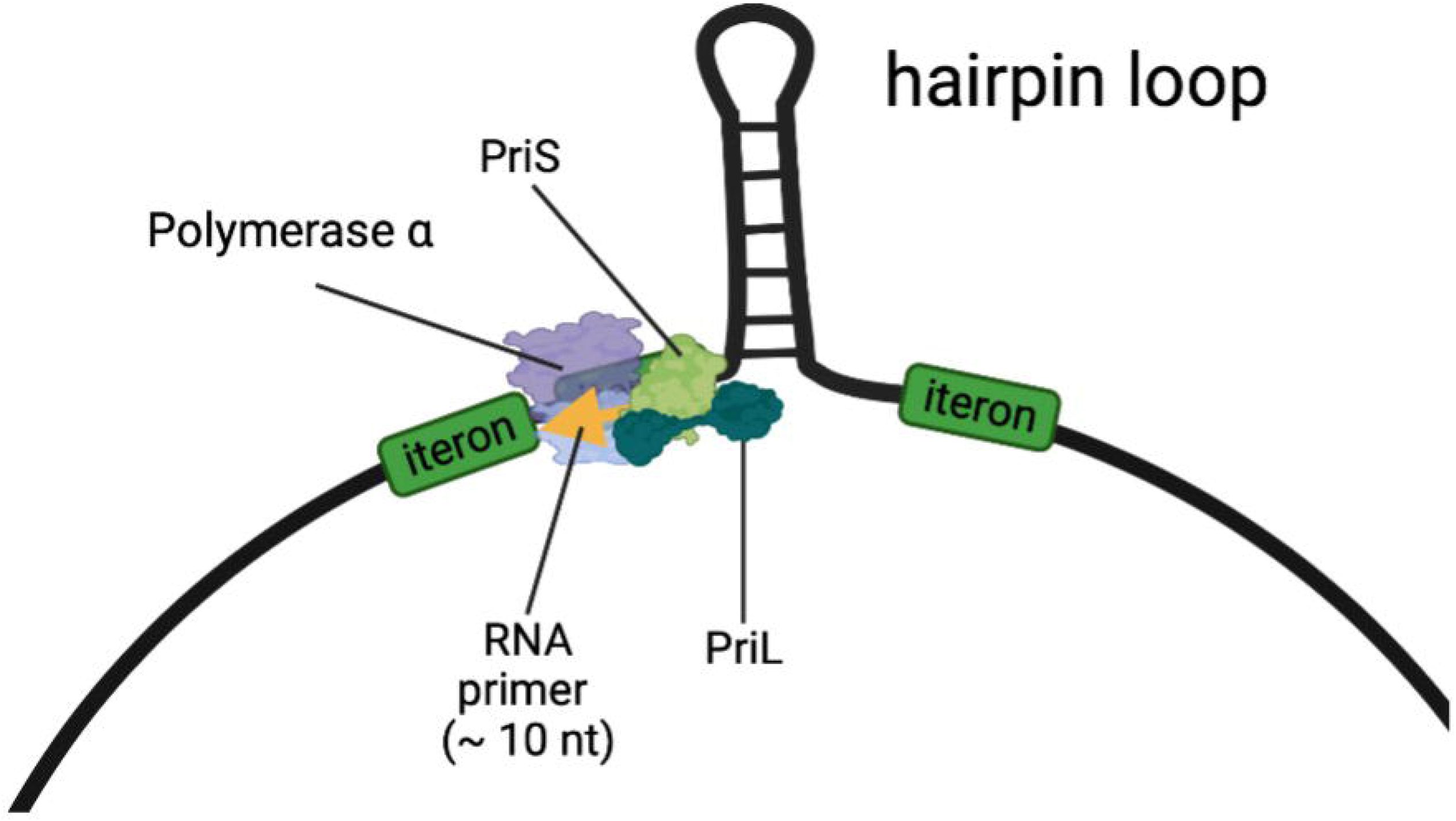
A model for the DNA Primase complex, consisting of PriL, PriS, and DNA polymerase α, producing an RNA primer near the hairpin loop of the ss-DNA of geminiviruses. DNA polymerase α consists of a catalytic subunit (p180) and an accessory subunit (p70). We hypothesize that in the resistant melon one or more mutations in PriL hamper the production or extension of the primer on the viral DNA, but still supporting the primer production in DNA replication forks of the host plant.

Mechanistically, it is yet still unclear how the changes detected in melon *PriL* specifically would inhibit its functionality towards the virus. The PriL protein comprises of two domains, an *N-*terminal (PriL_NTD_) and a *C-*terminal domain (PriL_CTD_) with a long, flexible linker of 18 residues in between (Baranovskiy et al., 2016). PriL_CTD_ is attached to the eukaryotic DNA of the lagging strand, supporting PriS with the initiation of the primer synthesis, while the PriL_NTD_ is attached to PriS which is elongating the primer by adding nucleotides (Baranovskiy et al., 2016). It is possible that ToLCNDV ss-DNA interacts with the PriL close to the *C-* terminus of the protein as this is the part that has DNA binding abilities and is unique in higher eukaryotes (Holt et al. 2018, Weiner et al., 2007, Klinge et al., 2007, Holzer et al., 2017). Mutations in this part could inhibit protein-viral DNA affinity and subsequently the efficacy of priming for viral DNA replication. However, the Tyr4His mutation in melon that associates with the resistance to the virus is located in the *N*-terminal of the protein (position 4). Mutations in the *N-*terminal could affect the interaction of the protein with the PriS or the Pol α (Baranovskiy et al., 2015), and in case of the latter abrogate extension of the RNA primer by Pol α. Alternatively, binding of other virulence factors (Fondong et al., 2013) of ToLCNDV to the N-terminal domain of PriL (PriL_NTD_) could be prevented and affect the functional activity and integrity of the primase/elongation complex. Considering that at this very first step in viral DNA replication, viral genome transcription has not yet occurred, and viral proteins thus are not synthesized, making the latter explanation less likely.

To understand the possible implications of PriL mutations for the discrimination between melon and geminivirus DNA replication, the model of the melon’s primase initiation complex was constructed. The model shows that the 3’-end of template DNA protrudes toward the open space at the junction of PriS and PriL_NTD_ (Figure 3). The major difference between the DNA templates of melon and geminivirus is a conserved 11 bp long stem-loop structure in a geminivirus circular ssDNA. Indeed, the space at the junction of PriS and PriL_NTD_ is a potential site for the docking of the stem-loop upon initiation of primer synthesis on the geminivirus circular DNA. Intriguingly, the Tyr4 and Pro125 are close to this area and their mutations may affect the primase interaction with the stem-loop, causing disruption of a delicate initiation process. The effect could be direct, by disrupting the local docking surface by Pro125Leu mutation or the primase interaction with the stem-loop by the Tyr4His mutation. Alternatively, the other viral or melon protein may facilitate the recruitment and holding of geminivirus stem-loop, and the Tyr4His and Pro125Leu mutations disrupt primase interaction with that accessory protein. Further studies are required to discriminate between these mechanisms.

**Figure 3.**
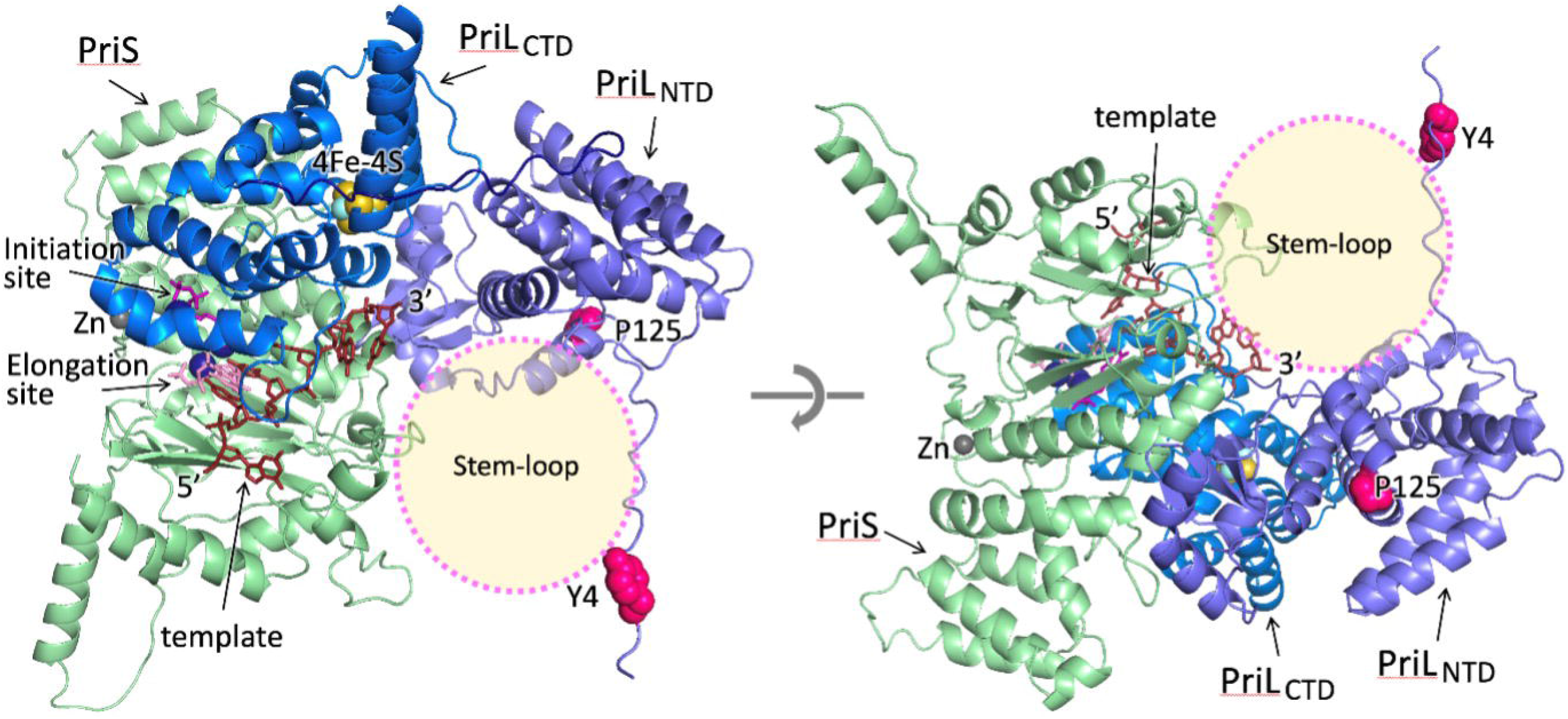
The model of the melon primase initiation complex. For clarity, two different views from opposite directions are shown. The potential stem-loop docking site is indicated by a transparent circle. The two amino acid substitutions in PriL in the resistant melon (Tyr4 and Pro125) are shown as circles in magenta.

## Conclusions

The results presented here provide supporting evidence for a role of *PriL* as an S-gene for geminiviruses. This evidence is based on sequence data analysis of the resistant melon accession K18 with the recessive QTL for resistance on ch11, and on DNA sequences of 100 other melon accessions and concomitant virus challenging assays to analyse their susceptibility or resistance to ToLCNDV. Additional support was obtained by silencing of *PriL* in *N. benthamiana* leading to reduced viral titers for three distinct geminivirus species. For almost 30 years the very first step in the conversion of ss-DNA to ds-DNA of geminiviruses has remained unclear. Based on the findings from this study, a role for the host DNA primase in the initiation of geminiviral DNA replication is postulated. How the Tyr4His and Pro125Leu amino acid substitutions within PriL of melon accession K18, carrying the Tomato leaf curl New Delhi virus (ToLCNDV) recessive resistance QTL in chromosome 11, is correlated to the resistance to ToLCNDV remains an experimental challenge for the future.

## Author Contributions

LS designed, conducted the research and wrote the manuscript, with inputs and supervision from HS, YB, RK and RV. MA contributed to the quantification of viral titers and gene expression for the VIGS experiment. JR, ME and CG contributed to the inoculation and disease scoring of melon accessions. DL provided the ToLCNDV resistant K18 melon for sequencing. TT and AB contributed to protein models for PriL.

## Supplementary Material

**Supplementary Table 1**. *PriL* alleles detected in various CGN accessions and the resulted phenotype after a challenge with ToLCNDV. The average ToLCNDV phenotypic disease score is given per accession and refers to a scale of 1 (highly resistant, no symptoms) to 9 (highly susceptible, severe symptoms). The resistant melon control genotype “Coliseo” exhibited a score of 1.5 whereas the susceptible melon control “Grand Riado” exhibited a score of 7.0. DI≤1.5 = Highly resistant, 1.5<DI≤3.0 = Intermediately resistant, 3.0<DI≤5 = Intermediately susceptible, 5<DI≤9 = Fully susceptible.

**Supplementary Table 2**. Primers used for qRT-PCR.

