## Supplementary tables for "*DNA primase large subunit* is an essential plant gene for geminiviruses, putatively priming viral ss-DNA replication"

| *PriL* allele | Phenotype (Number of accessions with the allele) | CGN accession (D.I score) |
| --- | --- | --- |
| AB | Highly resistant (1/1) | CGN140857 (1.0) |
| ABCD | Highly resistant (2/2) | K18, CGN140871 (1.0) |
| ACD | Highly resistant (4/5), Intermediately resistant (1/5) | CGN140839 (1.0), CGN24602 1), CGN24621 (1.0), CGN24624 (1.0), CGN140853 (2.9) |
| B | Intermediately resistant (1/1) | CGN140870 (1.8) |
| BCD | Intermediately susceptible (1/1) | CGN140868 (4.0) |
| BD | Intermediately resistant (1/1) | CGN140808 (1.8) |
| C | Fully susceptible (5/6), Intermediately susceptible (1/6) | CGN14805 (6.6), CGN14806 (4.8), CGN140858 (7.4), CGN140865 (7.8), CGN140866 (7.8), CGN140872 (7.8) |
| D | Fully susceptible (2/2) | CGN140807 (9), CGN140874 (6.4) |

### Supplementary table 1

### Supplementary table 2

| **Primer Set** | **Target** | **Forward** | **Reverse** |
| --- | --- | --- | --- |
| Nb*APR2 (Liu et al., 2012)* | *Solyc02g093150.2* | 5’-CATCAGTGTCGTTGCAGGTATT-3’ | 5’-GCAACTTCTTGGGTTTCCTCAT-3’ |
| Nb*PRIL1013* | *Niben101Scf00366g01013.1* | 5’- TCT GGA GGA TTT TGA GTT TTA TGC CAT- 3’ | 5’-TGCCTGCGGATGCCTCATATTTT- 3’ |
| Nb*PRIL1015* | *Niben101Scf01950g03015.1* | 5’-CCGCCATTTTCAGTATTCGACGAAATT-3’ | 5’-TACAGAACGATAGAGAGGCAGAGT-3’ |
| TolCNDV A (Simon et al., 2017) | ToLCNDV genomic component A | 5’-CATTATTGCACGAATTTCCG-3’ | 5’-ATCGTAGCCGACTGTGTCTG-3’ |
| TYLCV | TYLCV genomic sequence | 5’- TTCGTCTAGATATTCCCTATATGAGGAGGTA-3’ | 5’-GGCAAGCCCATTCAAATTAAAGG-3’ |
| BCTV | BCTV genomic sequence | 5’-TTGGGAAGAACAAATGGAGAACTCATTG-3’ | 5’-ATAGAGTAAAGCATTCTCCTTCACGTCTTC-3’ |
